# Auditory Steady-State Responses During And After A Stimulus: Cortical Sources, and the Influence of Attention and Musicality

**DOI:** 10.1101/2020.10.05.327189

**Authors:** Cassia Low Manting, Balazs Gulyas, Fredrik Ullén, Daniel Lundqvist

## Abstract

The auditory steady-state response (ASSR) is an oscillatory brain response generated by periodic auditory stimuli and originates mainly from the temporal auditory cortices. Recent data show that while the auditory cortices are indeed strongly activated by the stimulus when it is present (ON ASSR), the anatomical distribution of ASSR sources involves also parietal and frontal cortices, indicating that the ASSR is a more complex phenomenon than previously believed. Furthermore, while the ASSR typically continues to oscillate even after the stimulus has stopped (OFF ASSR), very little is known about the characteristics of the OFF ASSR and how it compares to the ON ASSR. Here, we assessed whether the OFF and ON ASSR powers are modulated by the stimulus properties (i.e. volume and pitch), selective attention, as well as individual musical sophistication. We also investigated the cortical source distribution of the OFF ASSR using a melody tracking task, in which attention was directed between uniquely amplitude-modulated melody streams that differed in pitch. The ON and OFF ASSRs were recorded with magnetoencephalography (MEG) on a group of participants varying from low to high degree of musical sophistication. Our results show that the OFF ASSR is distinctly different from the ON ASSR in nearly every aspect. While the ON ASSR was modulated by the stimulus properties and selective attention, the OFF ASSR was not influenced by any of these factors. Furthermore, while the ON ASSR was generated primarily from temporal sources, the OFF ASSR originated mainly from the frontal cortex. These findings challenge the notion that the OFF ASSR is merely a continuation of the ON ASSR. Rather, they suggest that the OFF ASSR is an internally-driven signal that develops from an initial sensory processing state (ON ASSR), with both types of ASSRs clearly differing in cortical representation and character. Furthermore, our results show that the ON ASSR power was enhanced by selective attention at cortical sources within each of the bilateral frontal, temporal, parietal and insular lobes. Finally, the ON ASSR proved sensitive to musicality, demonstrating positive correlations between musical sophistication and ASSR power, as well as with the degree of attentional ASSR modulation at the left and right parietal cortices. Taken together, these results show new aspects of the ASSR response, and demonstrate its usefulness as an effective tool for analysing how selective attention interacts with individual abilities in music perception.

## 1. Introduction

The auditory steady-state response (ASSR) is an oscillatory neural response that phase-locks to the presentation rate of a repeated auditory stimulus (e.g. a chain of clicks), or the modulation frequency (f_m_) of an amplitude- and/or frequency-modulated sound. It can be recorded using electro- and magnetoencephalography (EEG, MEG)^1-2^ and exhibits a maximum cortical power response at approximately 40 Hz in humans^1, 3^. The ASSR stabilizes at around 200 ms from when the stimulus begins, and continues to oscillate at a constant phase throughout the duration of the stimulus^4^. Since the ASSR’s oscillation frequency is equal to the stimulus’ f_m_, the ASSR power at this specific frequency can be extracted from MEG or EEG data in a straightforward manner using power spectral density (PSD) estimation techniques such as Fourier transformation. This offers a simple, efficient and precise method for separating neural responses to multiple simultaneously-presented auditory stimuli that are each modulated at a different f_m_^3, 5^. Hence, the ASSR has found itself useful in many applications ranging from hearing assessments^6-8^ to attention research^9-12^ and schizophrenia^13-14^.

However, questions such as what brain regions contribute to generating an ASSR, how an ASSR is influenced by cognitive processes and individual auditory expertise, or how the ASSR changes during and after a stimulus have not been fully explored. One popular theory regarding how the ASSR is generated suggests that the ASSR is a result of phase-locking between the envelope of the ongoing auditory stimulus and neuronal oscillations (hereafter referred to as the “ON ASSR”). This theory is dubbed the ‘Entrainment Hypothesis’, and is considered to give rise also to an extended “ringing” of the ASSR even after stimulus removal^4, 15-16^. While the presence of such post-stimulus ringing (hereafter referred to as the “OFF ASSR”) has been observed experimentally, the response characteristics and its underlying sources have not yet been examined. Given that the dominating view on cortical ASSR sources has been that the ASSR predominantly stem from temporal auditory cortices, the lack of research on the OFF ASSR may possibly reflect that researchers have assumed that it is a remnant activity leftover from a decaying ASSR with the same neural origins, and therefore should exhibit the same properties as the ASSR when the stimulus was present (ON ASSR).

Although this assumption is not unreasonable, more evidence is needed to substantiate it and clarify the association between the OFF ASSR and ON ASSR. This is an important step to achieving a fuller picture of the ASSR phenomenon and how to effectively utilize it. Recent data indicates that the ASSR involves more extensive cortical sources than previously believed, engaging both parietal and frontal cortices in addition to the temporal auditory cortices^9^. It is however not yet known what sources are engaged during the OFF ASSR period, and whether these sources differ from the ON ASSR. Furthermore, although several publications have shown that the ASSR power is modulated by selective auditory attention^11-12^, less is known about how the different cortical sources contribute to this modulation, and virtually nothing is known regarding whether selective attention influences the OFF ASSR per se. Finally, it remains unknown as to whether the ASSR itself or its modulation by selective attention is influenced by auditory experience and training such as musical expertise. Previous research has demonstrated that the neural correlates of auditory information processing vary with musical expertise^17-18^, and that the ASSR may be affected by age-related experience factors^19^.

In this study, our primary objective is to characterize and compare the ON and OFF ASSR in terms of their sensitivity to (i) stimulus physical properties such as volume and carrier frequency (f_c_), (ii) selective auditory attention, (iii) musicality, as well as their respective (iv) cortical source distributions. As a secondary objective, we will also follow-up on our recent work which showed that the ASSR is enhanced by selective auditory attention to a single melody stream presented within a mixture of three^9^. In the present study, we will extend this finding further by localizing the cortical areas of ASSR attentional modulation via statistical testing for each of the frontal, parietal, temporal and insular lobes. A priori, we hypothesize, based on our previous findings^9^,that all of these areas exhibit ASSR attentional enhancement. In addition, we will also explore the relationship between ASSR attentional modulation and musicality at each of these locations. Using MEG, we recorded the ON and OFF ASSRs from 29 adults with varying degrees of musical sophistication during a melody tracking task in which selective attention was shifted between different amplitude-modulated melody streams. Alongside sensor-space analyses, our source analysis adopted a minimum-norm estimate^20^ (MNE) distributed modelling approach to produce individual source estimates of these ASSRs.

## 2. Materials and Methods

### 2.1 Participants

A total of 29 participants with normal hearing volunteered to take part in the experiment (age 18 – 49 years, mean age = 28.6, SD = 6.2; 9 female; 2 left-handed). The experiment was approved by the Regional Ethics Review Board in Stockholm (Dnr: 2017/998-31/2). Both written and oral informed consent were obtained from all participants prior to the experiment. All participants received a monetary compensation of SEK 600 (∼ EUR 60).

### 2.2 The Goldsmith Musical Sophistication Index(Gold-MSI) self-report questionnaire

A subset of the Gold-MSI self-report questionnaire (v1.0)^21-22^ containing 22 questions was used to estimate each participant’s level of musical sophistication. These include all questions from the musical training and singing abilities subscales, selected questions from the perceptual abilities and active engagement subscale, but no questions from the emotions subscale. The MSI quantifies a participant’s level of musical skills, engagement and behaviour in multiple facets, and is ideal for testing amongst a general population that includes both musicians and non-musicians. Across all 29 participants, we obtained MSI scores ranging from 40 - 132, out of a maximum score of 154. A copy of the questionnaire used for this study can be found in the Supplementary Information (S1).

### 2.3 Experimental Task: Melody Development Tracking task

Participants performed the Melody Development Tracking (MDT) task according to the protocols in our previous study^9^, with the slight difference that a wider pitch separation was employed between the three frequency-tagged melody streams. Participants were presented with three melody streams of different pitch (i.e. f_c_) ranges, i.e. the Bottom voice, Middle voice, and Top voice. All three voices consisted of even (isochronous) melodic streams, but the three voices were slightly phase shifted relative to one another, so that the tones of each voice occurred separately in time as illustrated in Figure 1. The order of the tones was either Bottom-Middle-Top or the reverse, with the order balanced across trials. Before each melody began, participants were instructed to direct their attention exclusively to either the Bottom voice or the Top voice. When the melody stopped at a random time point, participants were asked to report the latest direction of pitch change between the two most recent notes in the attended voice (either *falling, rising* or *constant* pitch) with a button press (see Fig. 1). In total, 28 such responses were collected over approximately 15 minutes of MEG recording time for each participant.

**Figure 1.**
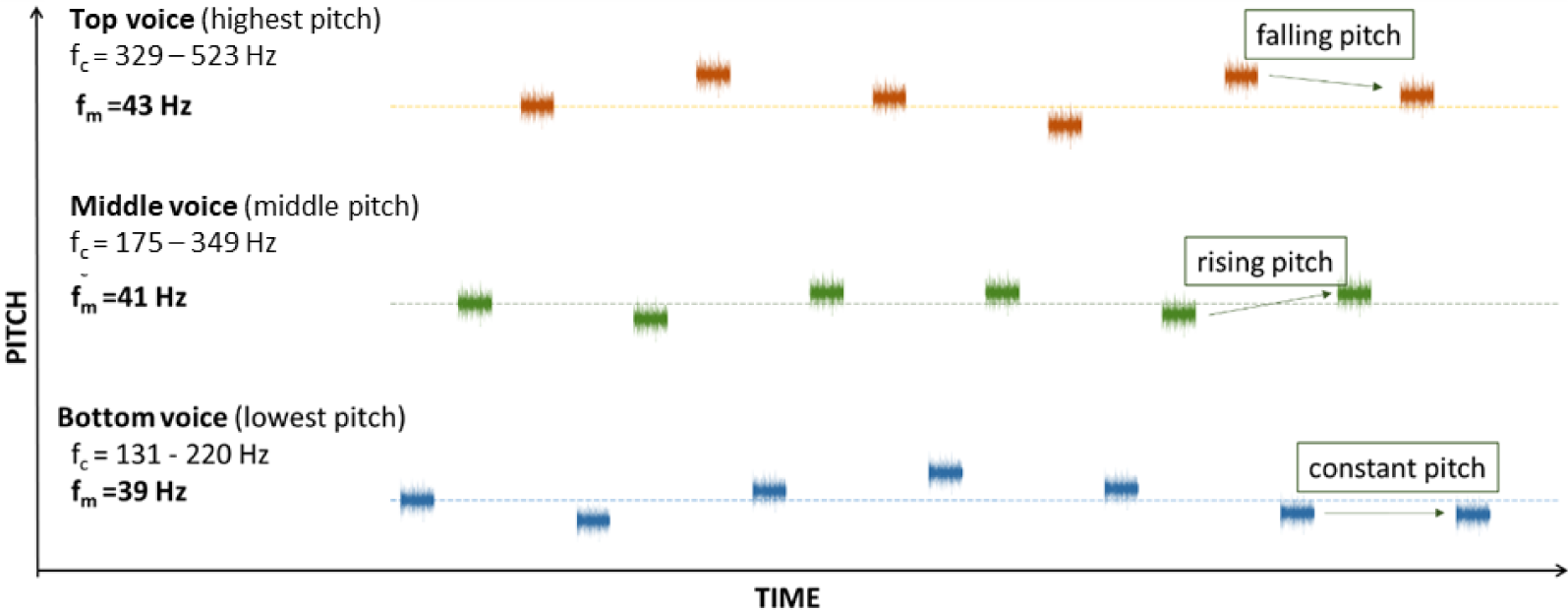
The Melody Development Tracking (MDT) task. Participants listened to three melody streams while attending to either the Bottom voice or the Top voice following an auditory cue. When the melody stopped, participants reported the latest direction of pitch change between the two most recent notes in the attended voice (i.e. falling, rising or constant pitch as illustrated). The three melody streams were presented separately in time, starting from Bottom to Top (as shown in figure) or its reverse. The respective f_c_ (pitch) range and f_m_ of each stream are indicated above.

### 2.4 Stimuli

Each of the three voices consists of a stream of 750 ms long sinusoidal tones of f_c_ between 131 – 329 Hz generated using the Ableton Live 9 software (Berlin, Germany). The tones that make up each voice stream were randomly-selected from the C major harmonic scale, allowing for repetition but limited to notes within their respective f_c_ range namely, the Bottom voice: 131 - 220 Hz, Middle voice 175 – 349 Hz, and Top voice 329 – 523 Hz. The minimum pitch difference between voices was 3 semitones. At the onset and offset of each tone, we introduced a 25 ms amplitude fade-in and fade-out to avoid audible compression clicks. These tones were then amplitude-modulated sinusoidally in Ableton Live 9 using f_m_ at 39 (Bottom voice), 41 (Middle voice), and 43 (Top voice) Hz, and a modulation depth of 100% to achieve maximum ASSR power^8^. Each tone is followed by 250 ms of silence before the next tone was played. The duration of melody presentation was randomized to be between 9 – 30 seconds long to reduce predictability of the stop point and thereby maintain high attention throughout the melody. At the stop point, the direction of pitch change was unique to each voice. The relative volume of each voice was adjusted to account for differences in subjective loudness for different frequency ranges^32^. The respective settings for the Bottom, Middle and Top voices were 0 dB, −6 dB and −10 dB, resulting in their raw volume decreasing in the same order. The stimulus was presented identically via ear tubes to both ears with the final average raw volume calibrated to 75 dB SPL per ear, subjected to individual comfort level, using a soundmeter (Type 2235, Brüel & Kjær, Nærum, Denmark).

### 2.5 Behavioural data analysis

To assess response accuracy in the MDT task, mean task performance scores (number of correct responses out of 28 total responses) were calculated across all conditions separately for each participant.

### 2.6 Data Acquisition

MEG measurements were carried out using a 306-channel whole-scalp neuromagnetometer system (Elekta TRIUXTM, Elekta Neuromag Oy, Helsinki, Finland). Data was recorded at a 1 kHz sampling rate, on-line bandpass filtered between 0.1- 330 Hz and stored for off-line analysis. Horizontal eye-movements and eye-blinks were monitored using horizontal and vertical bipolar electrooculography electrodes. Cardiac activity was monitored with bipolar electrocardiography electrodes attached below the left and right clavicle. Internal active shielding was active during MEG recordings to suppress electromagnetic artefacts from the surrounding environment. In preparation for the MEG measurement, each participant’s head shape was digitized using a Polhemus FASTRAK system. The participant’s head position and head movement were monitored during MEG recordings using head-position indicator (HPI) coils. Anatomical magnetic resonance images (MRIs) were acquired using hi-res Sagittal T1 weighted 3D IR-SPGR (inversion recovery spoiled gradient echo) images by a GE MR750 3 Tesla scanner, with the following pulse sequence parameters: 1 mm isotropic resolution, FoV 240 × 240 mm, acquisition matrix: 240 × 240, 180 slices 1 mm thick, bandwidth per pixel=347 Hz/pixel, Flip Angle=12 degrees, TI=400 ms, TE=2.4 ms, TR=5.5 ms resulting in a TR per slice of 1390 ms.

### 2.7 Data Processing

The acquired MEG data was pre-processed using MaxFilter (-v2.2)^33-34^, and subsequently analysed and processed using the Fieldtrip toolbox^35^ in MATLAB (Version 2016a, Mathworks Inc., Natick, MA), as well as the MNE-Python software^36^. Cortical reconstruction and volumetric segmentation of all participants’ MRI was performed with the FreeSurfer image analysis suite^37^.

#### 2.7.1 Pre-Processing

MEG data was MaxFiltered^23-24^ by applying temporal signal space separation (tSSS) to suppress artefacts from outside the MEG helmet and to compensate for head movement during recordings, before being transformed to a default head position. The tSSS had a buffer length of 10 s and a cut-off correlation coefficient of 0.98. The continuous MEG data were divided into 1 s-long epochs from stimulus onset (i.e. onset of each individual note). Epochs were then visually inspected for artefacts and outliers with high variance were rejected using *ft_rejectvisual*^35^. After cleaning, the remaining approximately 70 % of all epochs were kept for further analyses. The data was divided into six experimental conditions, consisting of epochs (∼100 per condition) for each of the three voices (Bottom, Middle, Top) under instructions to attend the Bottom voice or Top voice, respectively, i.e. i) Bottom voice × Attend Bottom (“Bottom-Attend”), ii) Bottom voice × Attend Top (“Bottom-Unattend”), iii) Top voice × Attend Top (“Top-Attend”), iv) Top voice × Attend Bottom (“Top-Unattend”), v) Middle voice × Attend Bottom, vi) Middle voice × Attend Top.

#### 2.7.2 Sensor-space analysis of MEG data

One participant was excluded due to lower-than-chance performance in the behavioural task, resulting in 28 participants for all sensor-space MEG analyses. We carried out sensor-space analysis on the cleaned MEG epochs to extract the ASSR power for each of the six conditions. Firstly, a 30 – 50 Hz bandpass filter was applied to the epochs which were then averaged per condition, resulting in the *timelocked ASSR*. The *timelocked ASSR* was then divided into two segments – an ON segment between 200 – 700 ms corresponding to the relatively stable part of the ASSR, and an OFF segment between 800 – 950 ms during which no auditory stimulus was present. Each of these segments were zero-padded to 1 s before applying a fast Fourier transform (hanning-tapered, frequency resolution = 1 Hz) to acquire separate power spectra for the ON and OFF ASSRs (Fig. 2). The ASSR power spectra were further averaged across all gradiometer sensors, after collapsing data from orthogonal planar gradiometers, to give the average gradiometer data per participant. Gradiometer sensors were selected for analysis as they are generally less noisy compared to magnetometers. From the ON and OFF power spectra, the maximum power across all frequencies was extracted accordingly for each of the six conditions to give the mean ASSR power per condition. This corresponds to the power at f_m_ for the ON condition (defined as 39, 41, and 43 Hz for the Bottom, Middle and Top voices respectively). As for the OFF ASSR, the maximum frequency value strays slightly from f_m_ as the decaying ASSR oscillation slows down, but is still within the expected value of around ∼40 Hz. For the analysis concerning ASSR power changes across voices (i.e. due to volume and f_c_), the Attend Bottom and Attend Top data were combined and averaged within each voice, resulting in the mean ASSR power per voice regardless of attention condition. Finally, the mean ASSR power was converted to the base 10 logarithmic scale [(lg(power)] to achieve a more normal distribution across participants for statistical analysis. Pearson’s correlation tests were used to investigate the relationship between lg(power) of the Middle voice (excluded in attention condition) and MSI for the ON and OFF ASSRs separately.

**Figure 2.**
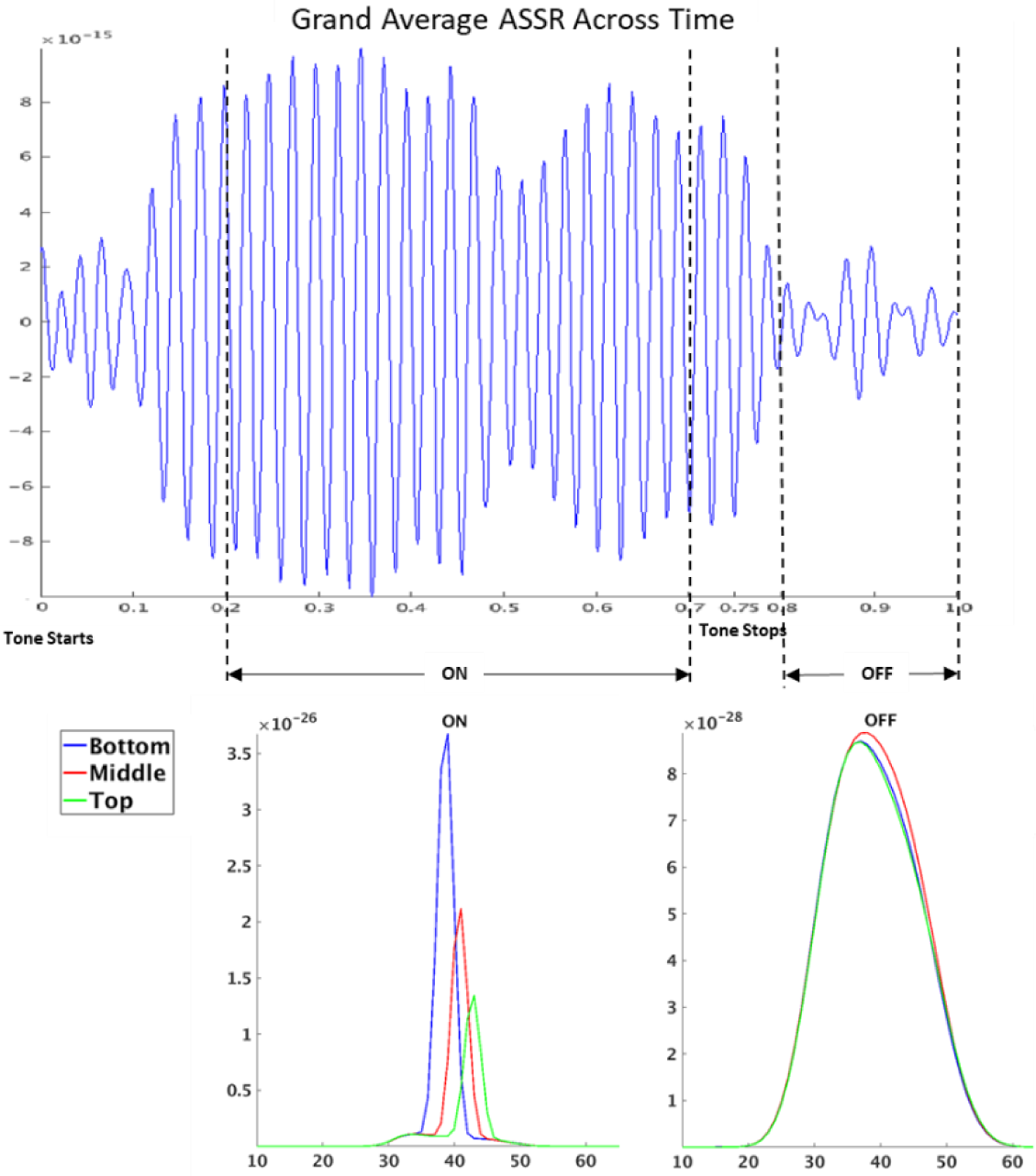
The ON and OFF ASSR power spectra. For each participant, the ON ASSR was obtained from a fast Fourier transform of the 200 – 700 ms time segment of the *timelocked ASSR* (top row), during which the tone stimulus was continuously present. Similarly, the OFF ASSR was similarly obtained from that of the 800 – 950 ms time segment, which occurred after the tone stimulus has stopped at 750 ms. The corresponding ON and OFF ASSR power spectra are shown in the bottom row. For the ON ASSR, the peak power occurred precisely at the modulation frequencies 39 (Bottom voice), 41 (Middle voice) and 43 Hz (Top voice). The OFF ASSR had wider, but clear, peaks at values lower than f_m_, albeit still close to 40 Hz, owing largely to the slowdown of the decaying ASSR. The across-subject grand average *timelocked ASSR* and frequency spectra are shown for illustrative purposes.

#### 2.7.3 Source-space analysis of MEG data

In addition to the participant excluded due to less-than-chance performance in the behavioural task, a second participant was excluded due to unsuccessful MRI collection, resulting in a final sample of 27 participants for the source-space MEG analyses. We used a minimum-norm estimate^20^ (MNE) distributed source model containing 20484 dipolar sources on the cortical surface to produce individual-specific anatomical layouts of the ASSR sources. These models were generated by inputting the ON and OFF sensor-space *timelocked ASSR* data into the MNE computation, before applying a Welch PSD estimation with zero-padding to 1 s (Hanning windowed, frequency resolution 1 Hz). Subsequently, the individual MNE solutions were morphed to a common fsaverage template.

#### 2.7.4 Source Pattern for ON vs OFF ASSR

We used the unattended Middle voice as a localizer to identify source distribution patterns of the ASSR across the cortex for the ON and OFF conditions. For each subject’s MNE solution, the power at each vertex was normalized by division over the maximum power across all 20484 vertices at 41 Hz, corresponding to the f_m_ of the Middle voice. This expresses the power of all sources as a fraction of the maximum power ASSR source, thereby correcting for individual power differences and allowing for subsequent averaging across subjects. The resultant MNE source pattern at 41 Hz represents the group overall distribution of the ASSR sources for ON and OFF.

#### 2.7.5 Location of the ON ASSR attentional enhancement and correlates with musical sophistication

Ten ROIs per hemisphere, across four lobes, were selected based on previous results^9^ showing that these ROIs contained spatial clusters which overall drive the ASSR attentional effect in a cluster permutation test. It should be noted, however, that the permutation test we applied only indicates an overall effect in the whole set of clusters, and not that the effect is present in each individual cluster. Here, we follow-up the question of the location of ASSR attentional enhancement, using a more confirmatory approach with the same ROIs as a priori target locations. The ten ROIs are the middle frontal gyrus, inferior frontal gyrus, orbital gyrus, precentral gyrus, superior temporal gyrus, middle temporal gyrus, posterior superior temporal sulcus, inferior parietal lobule, postcentral gyrus and insular gyrus. These ROIs are demarcated and classified into lobes according to the Brainnetome Atlas^25^ (see Fig. 5). Using the ON data, a MNE solution was computed for each Attend (Attend/Unattend) condition of the Bottom and Top voices (as the Middle voice was always unattended) per participant, and the ASSR power at f_m_ was extracted at 39 Hz and 43 Hz respectively. Next, all vertices within each ROI were averaged to give a median power value per ROI × Voice × Attend condition. The attentional modulation per ROI was expressed as a ratio between the Attend and Unattend conditions for each voice, and then averaged across the two voices. As a data reduction step, the ten ROIs were categorized into their respective lobes and averaged within the lobe (except for the insular lobe which only contained a single ROI), resulting in a total of eight datasets, each representing attentional modulation in one of the four lobes in the left or right hemisphere. As before, the Attend:Unattend ratio was converted to the base 10 logarithmic scale (lgAU) to achieve a more normal data distribution for statistical testing. The difference between lgAU and zero (since lgAU is equal to zero when the Attend:Unattend ratio is 1) was tested with one-tailed t-tests for positive attentional enhancement at a critical level of 0.05 with false discovery rate (FDR) correction over 8 tests using the Benjamini-Hochberg^26^ procedure. Additionally, Pearson’s correlations between lgAU and MSI were computed to quantify the relationship between ASSR attentional enhancement and musical sophistication at each lobe.

**Figure 3.**
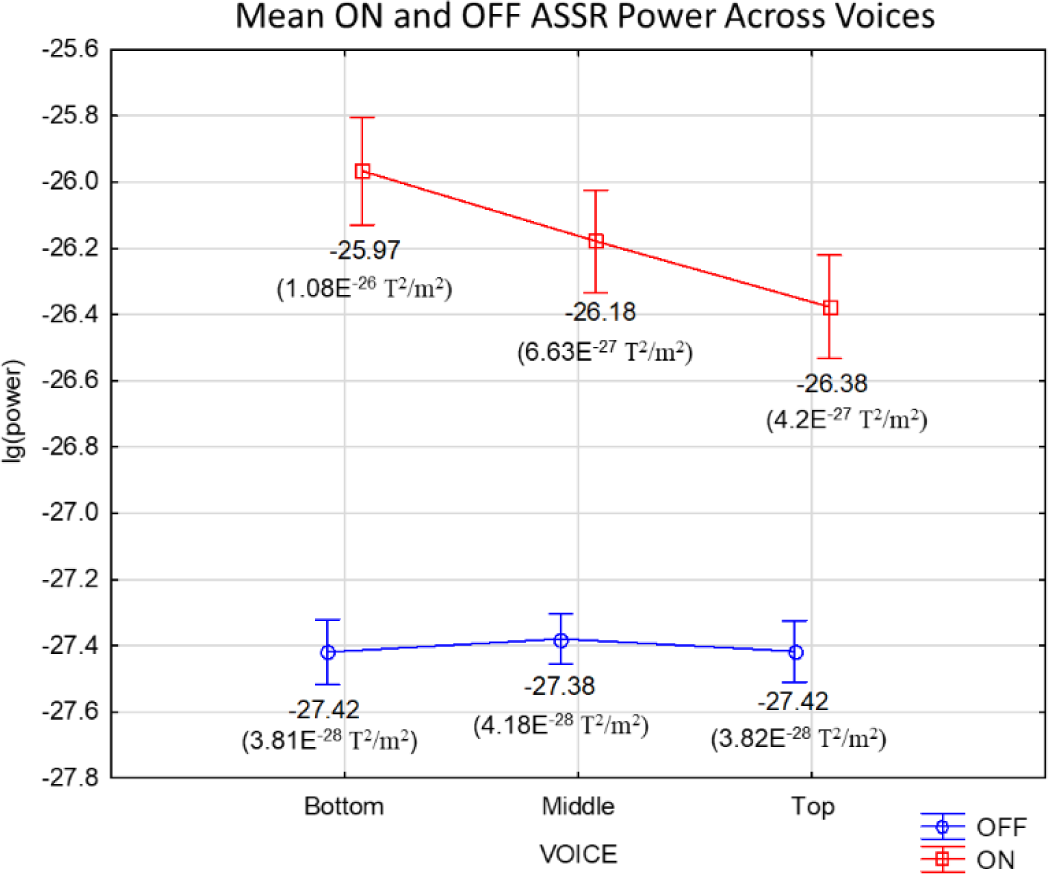
ON and OFF ASSR power across voices. Across-subject mean ON ASSR power decreases in the order Bottom-Middle-Top, as volume falls and f_c_ rises, whereas such a difference is hardly noticeable for the OFF ASSR. The lg(power) differences is significant (p<0.001) between all voices for ON and not significant between any voice for OFF in the post hoc Tukey tests (corrected). Furthermore, the ON ASSR displayed approximately 10 – 30 times stronger power than the OFF ASSR for all three voices. The corresponding ASSR powers in T^2^/m^2^, calculated directly from lg(power), are shown in parenthesis below the mean lg(power) values. Vertical bars denote 0.95 confidence intervals.

**Figure 4.**
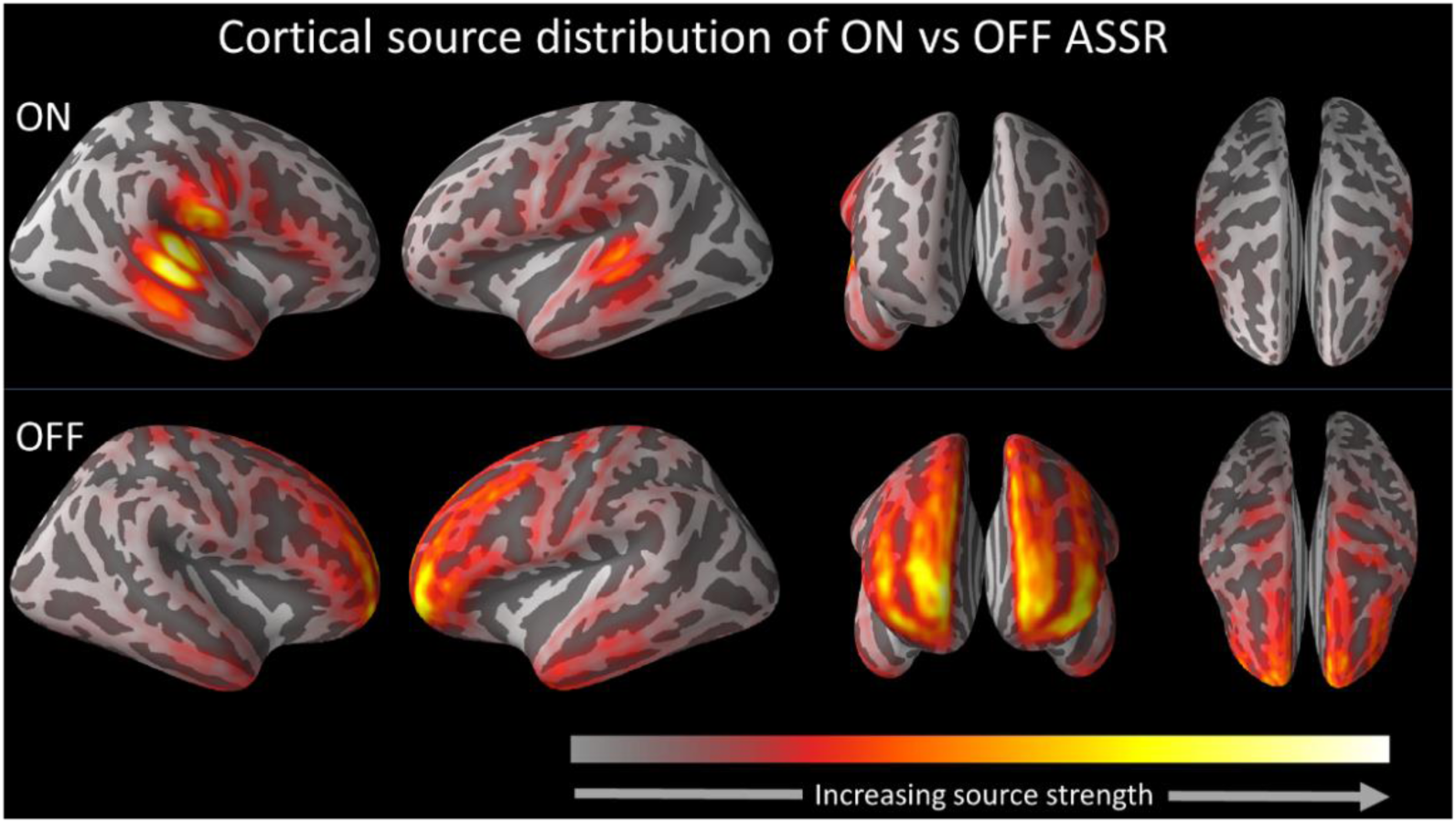
Comparison of the Middle voice cortical source distributions between ON (Top) and OFF (Bottom) ASSR. The across-subject grand average cortical source distribution maps at the peak Middle voice frequency are displayed. For the ON ASSR, the peak frequency is the f_m_ of 41 Hz, whereas for the OFF ASSR, it occurs slightly below f_m_ at 40 Hz as the oscillation slows down. The strongest ON ASSR sources were found in the primary and secondary auditory cortices within the temporal lobe. On the contrary, the strongest OFF ASSR sources resided mainly in the frontal cortices. Each map is normalized to its individual maximum source power (vertex-level) and scaled according to the colour bar shown. Orientation views from left to right: right lateral, left lateral, frontal, top.

**Figure 5.**
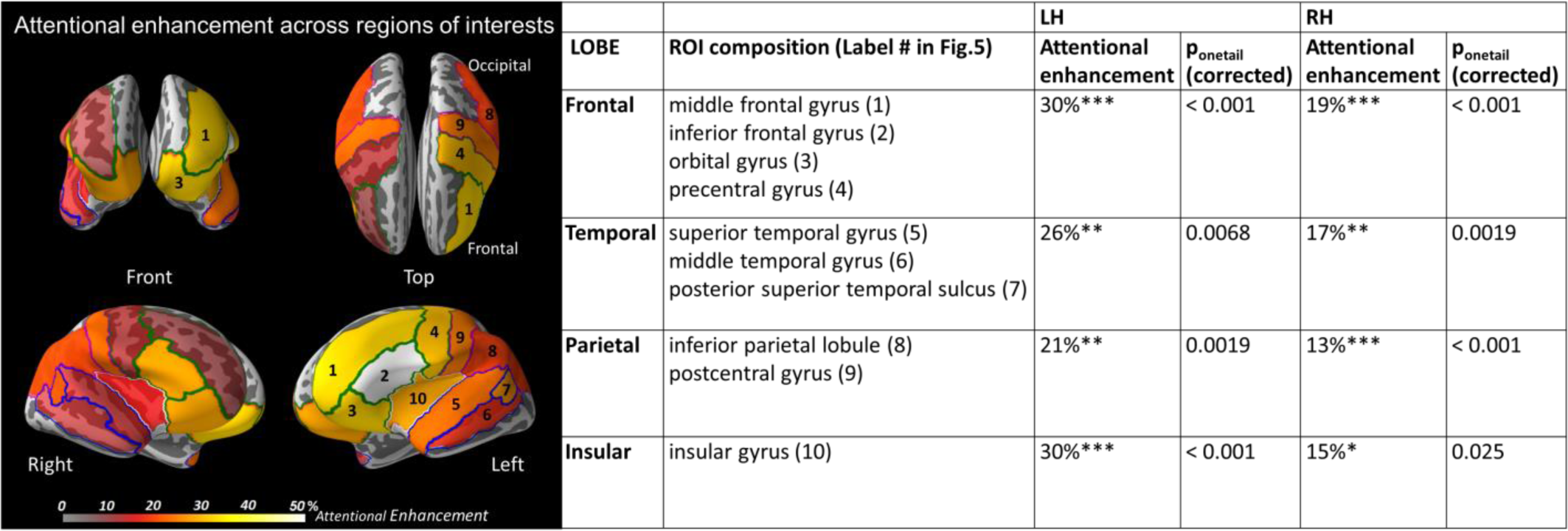
Across-subject grand average attentional enhancement across cortical lobes and regions of interests. (Left) The percentage increase due to selective attention is illustrated across the four lobes, with the constituent ROIs of each lobe demarcated in coloured lines (Green: Frontal, Blue: temporal, Magenta: parietal, White: Insular), and numbered according to column 2 (in parenthesis) of the table on the right. Colour bar is scaled to the percentage amount of attentional enhancement per ROI. (Right) The accompanying table displays the average attentional enhancement per lobe, back-calculated from lg(AU) values obtained from averaging across the ROIs that make up each lobe (columns 1 and 2). ALL FDR-corrected p-values from the one-tailed t-tests of lg(AU) values are reported in columns 4 and 6. p<0.001***, p<0.01**, p<0.05*

## 3. Results

### 3.1 MEG results

#### 3.1.1 Primary objective: Comparing the characteristics of the ON and OFF ASSR

For each subject, we computed the power spectra for the ON and OFF segments from the *ASSR_timelocked* data (see Fig. 2 for illustrative description) and compared them in terms of (i) voice differences (i.e. the stimulus volume and f_c_), (ii) selective attention, (iii) musicality (using the MSI), and (iv) cortical source distributions. Parts (i) – (iii) were obtained from the averaged sensor-level data, while part (iv) was addressed by MNE source analysis.

##### (i) ASSR power across voices

A repeated-measures ANOVA was used to test the differences in ASSR power across voices (Bottom/Middle/Top) and segments (ON/OFF). The analysis showed a significant main effect of segment (*F*[1, 27]= 357.5, *p* < 0.001), as well as voice (*F*[2, 54]= 62.2, *p* < 0.001). As can be seen in Figure 3, the ON segment (mean lg(power) across voices = −26.2) had ASSR responses of higher power than the OFF (mean lg[power] across voices = −27.4) segment. There was also a significant interaction between voice and segment (*F*[2, 54]= 75.3, *p* < 0.001, partial *η*^2^ = 0.736). Post-hoc Tukey tests revealed that there were significant differences between all three voices for the ON (p < 0.001, Bonferroni-corrected over 6 tests) but not for the OFF segment (minimum p value = 0.55, uncorrected). The decrease in ASSR power from Bottom to Middle to Top voice was expected since the sound volume was decreased in the same order across voices (as described under Methods)^*1*^. Figure 3 plots the lg(power) per voice for ON and OFF ASSRs, averaged across subjects. The corresponding ASSR powers in T^2^/m^2^ are computed and displayed in the figure.

##### (ii) ASSR power and selective attention

A separate repeated-measures ANOVA was used to test the modulation in ASSR power by selective attention (comparing Attend/Unattend conditions), voices (Bottom/Top) and segments (ON/OFF). As before, there was a significant main effect of segment (*F*[1, 27]= 280.7, *p* < 0.001), as well as voice (*F*[1, 27]= 118.2, *p* < 0.001). There was no main effect of attention (*F*[1, 27]= 10.8, *p* = 0.29). The analysis revealed a significant interaction between attention conditions and segment (*F*[1, 27] = 10.8, *p* = 0.0028, partial *η*^2^ = 0.285), and a post-hoc Tukey tests revealed significant differences between Attend and Unattend only in the ON time segment (*p* = 0.012) and not the OFF time segment (*p* = 0.58).

##### (iii) ASSR power and musical sophistication

Pearson’s correlation tests were run to quantify the relationship between ASSR [lg(power)] and MSI. For this analysis, only the Middle voice was tested for significant correlations with MSI. The results show a significant positive correlation between ASSR and MSI for the ON (*r* = 0.40, *p* = 0.036, *R*^2^ = 0.16; See Fig. 6d) but not the OFF ASSR (*p* = 0.58).

**Figure 6.**
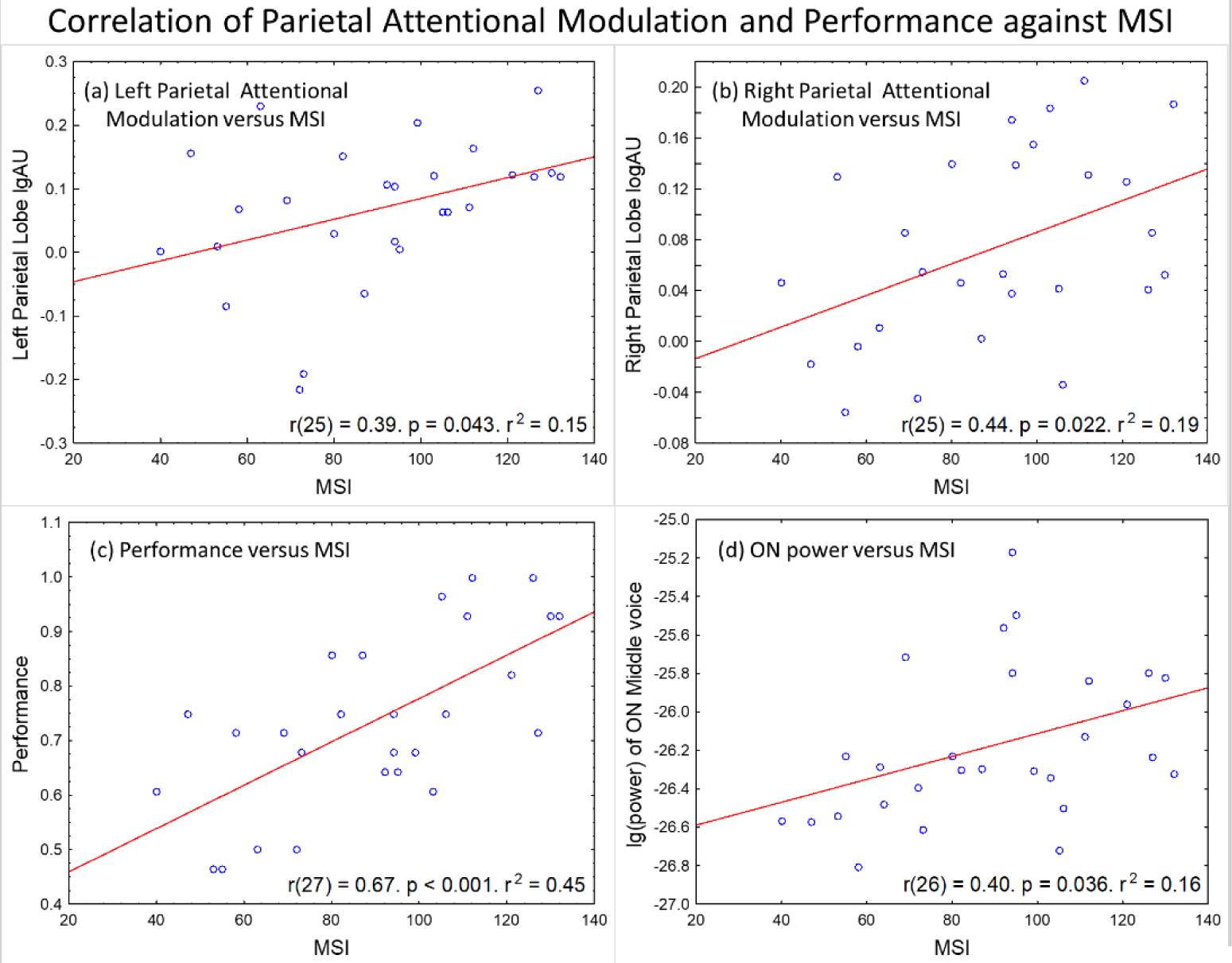
Correlation of left and right parietal attentional modulation and performance against MSI. Pearson’s r, R^2^ and p-values are indicated directly below each graph. (a & b) The attentional modulation across 27 participants correlates positively with MSI at the Left (*r*(25) = 0.39, *p* = 0.043, *R*^2^ = 0.15) and Right (*r*(25) = 0.44, *p* = 0.022, *R*^2^ = 0.19) parietal lobes. (c) Performance is positively correlated with MSI across all 29 participants (*r*(27) = 0.67, *p* < 0.001, *R*^2^ = 0.45). (d) ASSR power of the Middle voice is positively correlated with MSI for the ON segment (*r* = 0.40, *p* = 0.036, *R*^2^ = 0.16) across 28 participants.

##### (iv) Cortical ASSR source distribution

Using the unattended Middle voice as a localizer, we identified source distribution patterns of the ASSR across the cortex for the ON and OFF conditions. MNE source estimates show that the ON ASSR sources are localized primarily to the temporal auditory regions, whereas the OFF ASSR sources appear to be localized mainly to the frontal cortices. Figure 4 illustrates the cortical distribution of the ASSR signal for ON and OFF, averaged across all participants.

#### 3.1.2 Secondary objective: cortical source locations of ASSR attentional enhancement

##### (i) Attentional enhancement of ASSR power across lobes and hemispheres

Our second aim was to independently test if there was a significant enhancement of ASSR power due to selective attention in each of four lobes (frontal, temporal, parietal, insular) for two (left and right) hemispheres. Based on previous results from our group^9^, we hypothesized that the ASSR power in each of these areas would be significantly enhanced by selective attention. Figure 5 illustrates the across-subject grand average percentage attentional enhancement per ROI, categorized into their corresponding lobes by coloured borders (Green: Frontal, Blue: temporal, Magenta: parietal, White: Insular). Each ROI is labelled according to column 2 (in parenthesis) of the accompanying table on the right. The exact grand average attentional enhancement values corresponding to each ROI range from 9 – 50 % and can be found in the supplementary information (S1). The base 10 logarithmic of the Attend:Unattend ASSR power ratio [lg(AU)] was computed per lobe for each subject and used as a measure for attentional modulation. As hypothesized, the results from the t-tests yielded significant ASSR attention enhancement in all 8 lobes (FDR-corrected). The p values for each of these tests are displayed in the table on the right of Figure 5. The percentage enhancement values presented are back-calculated from the subject-median lg(AU) values to aid in interpretation. Alongside, Table 4 also shows the ROIs, with their corresponding label numbers in Figure 5, that comprise each lobe. A repeated-measures ANOVA using the lgADU values reported no main effect of hemisphere (*F*[1, 26]= 3.0, *p* = 0.096) or lobe (*F*[1, 26]= 1.6, *p* = 0.19), indicating that there was no significant difference in attentional enhancement between hemispheres or lobes.

##### (ii) The influence of musical sophistication on extent of ASSR attentional modulation

Finally, we investigated the relationship between musical sophistication and the degree of ASSR modulation from selective attention. By correlating individual MSI and lg(AU) values separately for each lobe and hemisphere, we found that the extent of ASSR attentional modulation in the parietal lobes (but not in the other lobes) was positively correlated with MSI: Left parietal lobe (*r*(25) = 0.39, *p* = 0.043, *R*^2^ = 0.15) and Right parietal lobe (*r*(25) = 0.44, *p* = 0.022, *R*^2^ = 0.19). With reference to Figure 6a and 6b, higher MSI was associated with a larger degree of ASSR attentional enhancement following selective attention. Figure 6 also depicts complementary correlations between MSI and performance (Fig. 6c), as well as between MSI and the ON ASSR power (Fig. 6d).

### 3.1 Behavioural results

As mentioned above, results from the MDT task showed that 28 out of 29 participants performed significantly above the chance level of 33% (M = 71 %, SD = 18.3 %; *t*(29) = 11.1, *p*_two-tailed_ < 0.001). These performance scores were positively correlated with MSI (*r*(27) = 0.67, *p* < 0.001, *R*^2^ = 0.45) (Fig. 6c), illustrating the sensitivity of the MDT task to individual level of musical sophistication.

## 4 Discussion

In this MEG study, we characterized the cortical ASSR that is generated during (ON) and after (OFF) the presentation of an amplitude-modulated stimulus using a task involving selective auditory attention to melody streams. We demonstrated that the ON and OFF ASSRs exhibit differences in their sensitivity to (i) stimulus volume and carrier frequency, (ii) selective attention and (iii) musical sophistication, as well as in their respective (iv) cortical source distributions. Furthermore, we statistically confirmed that the ASSR is enhanced by selective attention at cortical sources in each of the frontal, temporal, parietal and insular lobes. The following section discusses the key insights of this study.

### 4.1 ON and OFF ASSR have radically different characteristics

A novel and important finding in this study is that the OFF and ON ASSRs exhibited starkly different cortical source distributions, with the OFF sources mainly localized to the frontal cortex, while the strongest ON sources were localized to the temporal auditory cortices. While the presence of ASSR sources in the temporal^27-29^ and frontal^30-32^ cortices is in accordance with current literature and previous work from our group^9^, the discovery that frontal sources contributed to most of the OFF ASSR signal power was surprising and unprecedented. Moreover, the results also showed that the OFF and ON ASSRs exhibited different sensitivities to experimental factors and stimulus properties. The ON ASSR, on one hand, was modulated by several factors: It clearly decreased in power across voices from the Bottom to Top voice, reflecting the physical differences in volume and f_c_ between these voices. Furthermore, the ON ASSR power correlated positively with individual MSI, and significantly increased with selective attention.

On the other hand, the OFF ASSR, remained unaffected by the same factors, displaying very similar power levels across voices, and was neither affected by MSI nor selective attention. Collectively, these findings indicate that the ON and OFF ASSR reflect very different cortical processes, and that the OFF ASSR is not simply a ringing extension generated by the same sources that underlie the ON ASSR. While the ASSR itself reflects a neural representation of the periodic envelope of the stimulus, the pattern of results suggests that the brain progresses from a sensory processing and discrimination state (i.e. identifying and paying attention to a tone) driving the ON ASSR at the auditory cortices, to a different frontal-dominated state during which the periodic neural representation is still maintained (i.e. the OFF ASSR), but is not sensitive to external factors such as the stimulus’ properties or to the perception of and attention to it. To the best of our knowledge, this is the first time any study has characterized the OFF ASSR, and the inclusion of this novel OFF ASSR adds a new dimension to the existing literature and potential uses of the ASSR for research, especially in the neuroscientific understanding of higher executive processing.

### 4.2 Selective attention enhances ASSR in multiple cortical lobes

In agreement with our hypotheses, our results demonstrate a significant attentional enhancement of the ON ASSR in each of the frontal, temporal, parietal and insular lobes across both hemispheres. This enhancement was strongest in the frontal cortex, the established centre for attentional control^33-35^, with an average ASSR enhancement of 30 % and 19 % for the left and right hemispheres respectively. Compared to our previous work results^9^ which suggested that the ASSR attentional enhancement can reach up to 80 % in frontal regions^9^, the current findings yielded substantially lower values. We believe that the differences between these studies can be explained by the thresholds used in the previous analysis to exclude vertices with weak ASSR power, whereas in the current analysis, the thresholding step was omitted to ensure that all ROIs contained sufficient vertices to be averaged at a single subject level. With the inclusion of vertices with little or no ASSR activity during averaging in this study, it is reasonable to expect the computed ASSR powers and attentional modulation to be diluted and thus reduced.

When discussing the individual signal contribution from different cortical areas, a critical point to consider is of course the degree of independence of these sources. With MEG (and EEG), field spread is an unavoidable aspect of the MEG signal. We addressed this issue in depth with point spread simulations in our previous study^9^ demonstrating a clear independence from frontal and temporal sources. Similarly, the characteristics of the current results in Figure 5, suggest that the frontal sources are spatially separated, and thus likely to be independent from the parietal, temporal and insular sources. However, the ASSR sources in the latter 3 lobes often appear relatively close to one another, and in some cases even connected. It is then important to consider whether some of these sources arise from independent sources overlapping in space that cannot be differentiated by our source localization, or whether they are artefacts of signal spreading. Nonetheless, at the very least, the present results provide the first confirmatory evidence substantiating that selective attention enhances the ASSR power at the frontal cortex.

### 4.3 Musical sophistication is associated with an increase in ASSR power and attentional enhancement, especially in the parietal cortex

Our results also showed that that musical sophistication influences the ON ASSR, both the ASSR power itself, and the degree of its modulation from selective attention. Notably, the behavioural results showed a significant positive correlation between the MSI of participants and their performance scores in the MDT task (*r* = 0.67), demonstrating the musical sensitivity of the task. More importantly, our results show that the ASSR power correlates strongly with MSI scores (*r* = 0.40). These results are in agreement with previous studies^18, 36-37^ showing that neural processing of auditory stimuli are enhanced by musical abilities and experience, and may reflect an improvement in auditory skills owing to musical training. Additionally, we found a strong positive correlation between MSI and the degree of attentional enhancement in the left (*r* = 0.39) and right (*r* = 0.44) parietal cortices with no such relationships to attentional enhancement in temporal and frontal cortices. In earlier studies^38-40^, parietal regions have been shown to exhibit sensitivity to musical experience and ability. This has been attributed to the fact that these regions play important roles for musical skill learning and performance. These evidences further strengthen the belief that musical experience is related to selective listening ability, a phenomenon which can be reflected in recorded neural signals like the ASSR. Speculatively, even better correlations between ASSR and musical experience may be achieved by using more musically-relevant or challenging experimental tasks with greater specificity and sensitivity to individual musical experiences and listening ability in future studies. For example, one could use more natural auditory stimuli like tones from a particular instrument that the musician participant has been trained in.

### 4.4 Conclusions

To summarize, we demonstrated that ON and OFF ASSR are driven by different cortical sources ? the frontal cortex for the OFF ASSR, and the temporal cortex for the ON ASSR. Furthermore, while the ON ASSR power was influenced both by physical stimulus features, selective attention and musicality, the OFF ASSR was not modulated by any of these factors, suggesting that the ON and OFF ASSRs stem from distinct neural states. Importantly, we have also confirmed that the ON ASSR exhibits significant attentional enhancement in the bilateral frontal, temporal, parietal and insular lobes, thus confirming our initial hypothesis. Moreover, we found that the degree of attentional enhancement correlated positively with individual musical sophistication scores specifically at the left and right parietal cortices, areas that are commonly associated with musical training^38-40^. Taken together, these findings deepen our understanding of the ASSR, while demonstrating it as a suitable and effective tool for analyses of higher cognitive processes such as music perception and selective attention.

## Acknowledgments

Data for this study was collected at NatMEG, the National Facility for Magnetoencephalography (http://natmeg.se), Karolinska Institutet, Sweden. The NatMEG facility is supported by Knut and Alice Wallenberg Foundation (KAW2011.0207). This study was supported by the Swedish Foundation for Strategic Research (SBE 13-0115).

## Supplementary Information

### S1 Subset of MSI Questionnaire used in this study

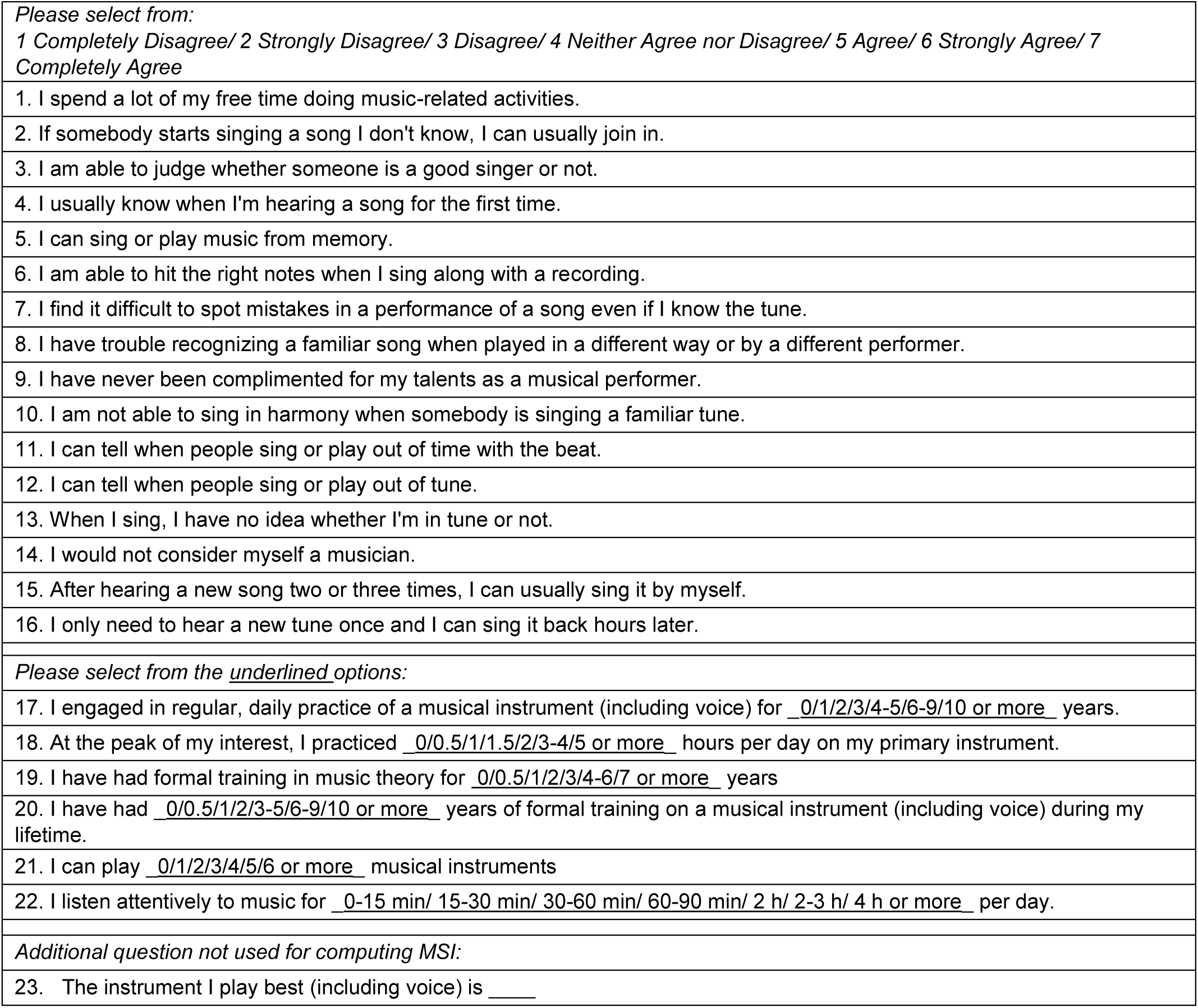

### S2 Table of across-subject grand ASSR attentional enhancement per ROI

| LOBE | ROI (Label # in Fig.5) | Average Attentional Enhancement |  |
| --- | --- | --- | --- |
|  |  | LH | RH |
| Frontal | middle frontal gyrus (1) | 31% | 9% |
|  | inferior frontal gyrus (2) | 50% | 27% |
|  | orbital gyrus (3) | 31% | 27% |
|  | precentral gyrus (4) | 29% | 11% |
| Temporal | superior temporal gyrus (5) | 22% | 12% |
|  | middle temporal gyrus (6) | 17% | 11% |
|  | posterior superior temporal sulcus (7) | 25% | 19% |
| Parietal | inferior parietal lobule (8) | 18% | 19% |
|  | postcentral gyrus (9) | 23% | 22% |
| Insular | insular gyrus (10) | 27% | 15% |

